# Clustering of Direct and Indirect DNA Binding Motifs of Human and Mouse Transcription Factors: X-TFBS Compendium from ChIP-seq

**DOI:** 10.1101/2025.11.09.687492

**Authors:** Alexei A. Sharov

## Abstract

Uncovering networks of gene expression regulation requires knowledge of specific DNA-binding motifs of transcription factors (TFs). Most TFs have multiple DNA motifs enriched in its ChIP-seq peak regions because of protein interaction and spatial correlation between TFs and cofactors. To capture both direct binding and indirect association of TFs with specific DNA locations, all-against-all relations are identified here between TFs and DNA-binding motifs by the reanalysis of 8027 human and 2820 mouse ChIP-seq experiments from GEO. DNA motifs were analyzed with CisFinder and then clustered using the new k-mean algorithm tailored for this kind of data. Additional clusters of motifs were found by filtering ChIP-seq peaks based on their location in promoters, enhancers, and repeat-depleted regions. The new X-TFBS compendium, which includes 1157 human and 536 mouse clusters bound by 459 orthologous human-mouse pairs of TFs, 1038 human-only, and 165 mouse-only TFs, is the largest among existing databases. Most orthologous TFs in human and mouse have nearly identical DNA-binding motifs. Clustering helps to annotate TF-binding motifs and evaluate interactions between TFs that are associated with the same motif. Visual comparison of large sets of DNA motifs is simplified by using sequence script instead of sequence logo.

## 1. Background

The goal of this paper is to explore binding of transcription factors (TFs) to specific DNA sequence motifs, which may regulate the expression of nearby genes in human and mouse cells. The most reliable method for finding DNA-binding motifs *in vivo* is ChIP-seq (Nakato and Shirahige 2016; Park 2009). But in contrast to *in vitro* methods, such as SELEX and protein binding microarrays, ChIP-seq captures both direct and indirect binding of TFs to DNA motifs. The latter is mediated by other TFs, cofactors, histones, RNA, and epigenetic modifications. These additional factors allow binding of individual TFs to multiple types of DNA motifs, and some of additional motifs may be functional. Also, ChIP-seq captures motifs of non-target TFs if they are spatially correlated with motifs of target TFs, which creates ambiguity in the interpretation of data, but also may provide valuable information about potential interactions of co-localized TFs. The effect of spatial correlation is excluded in the DNase I cleavage footprinting method (Neph et al. 2012; Vierstra et al. 2020).

To identify both direct and indirect binding of TFs to DNA motifs, Ambrosini et al. (Ambrosini et al. 2020) proposed an all-against-all approach, where each ChIP-seq experiment with a specific TF is evaluated against all possible motifs enriched in peaks of DNA reads, and applied this approach to published TF binding motifs. Objectives of Ambrosini et al. included (i) finding best overrepresented motifs for each TF independently of original motif annotation, (ii) distinguishing consistent differences in binding motifs of two TFs from random noise, (iii) removing redundancy in TF binding motifs, (iv) suggesting hypothetical mechanisms of in-vivo TF binding to sites with different motifs (e.g., in a different cell type or stimulation), and (v) discovery of cooperative binding of several TFs to the same motif. Similarity of two different position-weight matrixes was assessed using the Jackard index applied to two corresponding sets of matching sites found within a long random DNA sequence (Vorontsov et al. 2013). Another method to evaluate similarity of motifs is TOMTOM in the MEME Suite https://meme-suite.org/meme (Gupta et al. 2007).

These analytical approaches and statistical tools move the study of TF binding along a new path towards functional understanding of transcription regulation. However, this transition has just started and various obstacles and gaps still remain. First, available databases of DNA binding motifs, e.g., (Heinz et al. 2010; Lambert et al. 2018; Pratt et al. 2022; Vorontsov et al. 2024) do not include many ChIP-seq data in the GEO. Second, binding motifs are identified from all binding sites irrespective of their location in the genome relative to promoters, enhancers, and repeats. This strategy may appear not sufficiently effective for those binding motifs that are preferentially located either in promoters, or enhancers, or repeat regions. Third, the k-mean clustering algorithm has not been implemented for the analysis of DNA binding motifs. Instead, the hierarchical clustering algorithm with a fixed threshold similarity was used by Ambrosini et al. (2020), which is less efficient than k-mean method in cases where motif variance is not uniform across clusters. And fourth, visualizing and managing of large clusters of similar binding motifs requires new tools. Sequence logo (Schneider and Stephens 1990), which is the most frequently used visualization method, is applicable for comparison of a few motifs and cannot be used for visual comparison of hundreds or thousands of similar motifs. In addition, sequence logos cannot be easily copied, moved, and searched within standard software, such as MS Excel.

This study fills some of the gaps in our knowledge of TF binding motifs by systematic analysis of the majority of human and mouse ChIP-seq experiments deposited in the GEO database. I adopt the all-against-all approach (Ambrosini et al. 2020) for the analysis of TF binding motifs and focus on finding matching motifs in multiple independent ChIP-seq experiments. The greater is the number of experiments that yield matching motifs, the more reliable are these motifs. The number of DNA motif clusters discovered in this study approximately doubles the number of motif clusters in existing databases. A new ‘sequence script’ method is developed and used for visualization of large sets of aligned motifs. Sharing the same DNA-binding motif by several TFs often indicates potential functional interaction between them, which can prompt the discovery of new molecular signaling pathways.

## 2. Methods

### 2.1. Finding DNA-binding motifs of transcription factors from ChIP-seq data

This project is based on the reanalysis of 8027 human and 2908 mouse ChIP-seq experiments from GEO (Supplementary file S1). Preference is given to data with genome coordinates of ChIP-seq peaks (98% of data). Alternatively, peaks were identified from files in bigwig, wig, bed, or eland formats. Peak coordinates were converted to hg38 human and mm10 mouse genome versions using the UCSC Liftover tool (https://genome.ucsc.edu/cgi-bin/hgLiftOver). DNA-binding motifs were identified using CisFinder (Sharov and Ko 2009), in particular, its stand-alone tools patternFind, which generates elementary motifs based on the frequency of all 8-bp “words” (continuous and with gaps) in the DNA sequence, and patternCluster, which merges elementary motifs into a longer composite motif^1^. Motifs were found from comparison of 200-bp (or 120-bp) genome segments centered at ChIP-seq peaks with two control 150-bp genome fragments separated by 500 bp from the peak fragment on both sides. Only one motif instance was counted per peak to avoid counting multiple instances in repeat regions. Human motifs were identified first by using 200-bp long DNA segments in 5,000 top-scored peaks, and second, using 120-bp long DNA segments in 10,000 top-scored peaks. The second method was designed to increase the spatial resolution of analysis, but it did not provide sufficient advantage. Because these methods generated very similar results, only the first method was used to find DNA motifs in mouse. For each ChIP-seq sample, not more than 5 top-score motifs were selected for further analysis; motifs supported by less than 30 ChIP-seq peaks were not processed further. The frequency of each motif was estimated by searching for motif instances in ChIP-seq peak genome segments with CisFinder stand-alone tool patternScan with false positives rate of 1 per 100 Kb of random DNA sequence length.

A large portion of functional DNA-binding motifs of TFs are located in evolutionary-conserved regions that include promoters and enhancers; and thus, the initial approach was to filter out those ChIP-seq peaks, that contained more than 70% of repeats based on repeatmasker output downloaded from UCSC (Navarro Gonzalez et al. 2020), and motifs identified in repeats were discarded. However, motifs of some TFs, such as zinc finger proteins, are more abundant in repeat regions (Imbeault et al. 2017); and thus, motif search was repeated without filtering peaks and avoiding repeat regions. These two approaches appear to complement each other, because each of them yields motifs that are not detected with another method. A natural extension of this approach was to filter ChIP-seq peaks based on their location in promoter regions (−300 to 300 bp relative to TSS^2^), or in previously identified enhancers (Gao and Qian 2020), and more new motifs were found in this way. The compendium of DNA-binding motifs presented in this paper combines results of all these computational methods.

### 2.2. Clustering of DNA-binding motifs with a modified k-mean method

To cluster TF-binding motifs, I developed a k-mean clustering program optimized for fast processing of large sets of data (https://github.com/AlexeiSharovBaltimore/CisFinder; file: kmean_motif.c). The distance/dissimilarity between motifs is estimated as *d* = 100·(1 - *r*), where *r* is a Pearson correlation of transformed position-weight matrix (PWM) data: *p*’ = 0.75 + 3·(*p* – 0.75) if probability *p* > 0.75, and *p*’ = *p* otherwise. This simple transformation increases the contribution of high-frequency nucleotides to the distance measure. The distance is then minimized for direct and reverse orientations and position shifts of compared motifs relative to each other. Position shifts are limited to ensure the minimum overlap length of 6 nucleotides. Frequency of overhanging nucleotides is set to the overall average value for each nucleotide. Clusters are initiated with a set of randomly selected single motifs. At each iteration, closely located clusters with distance between centers <3 are merged, and then, created cluster vacancies are filled with randomly selected member motifs located far from their cluster centers. The iteration stops when the overall square distance to cluster centers is minimized, or if cluster membership remains the same. Although the distance measure is not Euclidian, it supports the conversion of the k-mean iterative process.

Each cluster often includes multiple similar motifs obtained from the same ChIP-seq sample. Thus, to reduce redundancy, I removed redundant motifs from the same sample in a given cluster except for only one or two most informative ones and/or originated from non-repetitive genomic regions, promoters, and enhancers. The latter condition is important to preserve evolutionary conserved motifs, which are more likely to be functional in regulating gene expression than non-conserved motifs.

For each motif cluster, an integrated motif was built based on 3 top motifs in the cluster, which were selected manually using the following criteria: sufficient motif length, number of motif instances in ChIP-seq peaks, and consistency between these 3 motifs. In each motif position, nucleotide frequencies were averaged with weights equal to number of motif instances. If two motifs have the same dominating nucleotide in a given position that is different from the third motif, then the third motif was not used in estimation of frequencies in that position.

Traditionally, DNA-binding motifs are attributed to a single TF targeted by the antibody in ChIP-seq. However, the all-against-all approach requires a more flexible naming convention for DNA-binding motifs. Here, the name of a motif cluster starts with the name of the main TF that presumably binds it directly based on available data (HOMER, HOCOMOCO, dbTFBS). Some names of TFs are abbreviated to represent a group of similar TFs that bind the same motif (e.g., ATF, SOX, FOX, KLF). However, names of C2H2 zinc-finger proteins (e.g., ZNF, ZFP) are not abbreviated because most of them have unique motifs as a consequence of their faster evolution in comparison to other TFs (Lambert et al. 2019). If the directly binding TF is unknown or not present in the set of analyzed ChIP-seq samples, then the first TF in the motif name is selected based on the high frequency of its association with the motif. Clusters of motifs are ordered alphabetically, and this order helps to compare motifs of closely-related TFs whose names differ only by ending characters. To facilitate such comparison, I added abbreviated names of TF classes in front of names of some TFs: bHLH – for basic helix-loop-helix, bZIP – for basic leucine zipper, HOX – for homeo domain, NR – for nuclear receptor, and ETS – for ETS-related factor. These prefixes are added not globally, but solely for the convenience of motif sorting. In particular, they are not applied to TFs with distinct motifs, such as IRF and POU factors in the homeo domain class.

The name of the main TF is followed by the names of other TFs that are consistently represented in the cluster and associated with the same motif. To distinguish multiple motif clusters associated with the same TF, I used additional tags in cluster names, such as “altA”, “altB”, for alternative motifs, “pal” for palindromes, and “short” for short versions of the motif. Names of motif clusters that are detected predominantly in promoters, enhancers, or repeats include tags “promoter”, “enhancer”, or “repeat”, respectively. Motifs supported by a single ChIP-seq experiment are marked as “weak”, indicating that they require further confirmation. The exception is given to alternative motifs that resemble a well-confirmed main motif and to motifs supported by experiments in both human and mouse. If a cluster of motifs is strongly enriched in specific transposable elements; then the name of repeat elements (repeatMasker) is included in the name of the cluster. Motif clusters that have no dominating TFs are considered non-specific and named “NONSPEC” followed by a number. If such a motif resembles a known TF-specific motif, then it is marked as, for example, “MYC-like” or “REST-like”.

### 2.3. New method for motif visualization: sequence script

Sequence script method combines upper-case, lower-case characters, and the underscore character; and it is designed to emphasize the core portion of DNA-binding motifs, and allows easy comparison, copying, pasting, and searching in large sets of motifs. A sequence script is assembled based on the position-frequency matrix (PFM) as follows. In each position of the motif, nucleotides are sorted by decreasing frequency (or probability). If the frequency of the dominating nucleotide (rank-1) is ≥ 2-fold of the probability of the rank-2 nucleotide, then the upper-case dominating nucleotide is entered in the sequence script (Fig. 1, strong dominance). If this condition is not met, but the probability of the rank-1 nucleotide is ≥ 1.3-fold of the probability of the rank-2 nucleotide, then the lower-case dominating nucleotide is entered in the sequence script (Fig. 1, weak dominance). If the previous condition is not met, but the probability of the rank-2 nucleotide is ≥ 2-fold of the probability of the rank-3 nucleotide, then a lower-case letter representing the pair of first two nucleotides (w, s, r, y, m, and k for nucleotides AT, CG, AG, CT, AC, and GT, respectively) is entered in the sequence script (Fig. 1, pair prevalence). Finally, if none of these conditions are met, then the underscore symbol “_” is entered, which means no prevalence. If the underscore symbol(s) appears at the start or end of the sequence scrip then it is removed (trimmed). To align multiple sequence scripts it is necessary to use equal width (monospaced) fonts, such as Courier, Consolas, or Monaco. The best software to compare sequence strings is Excel or Google Sheets because, first, it allows using space characters at the start to move a string left and right to align it with other strings, and second, it allows wild card characters for finding motifs. For example, the JUN/FOS binding motifs can be searched as “TGA?TCA”, where the question mark is a wild card representing any character in a sequence string. In supplementary tables, all motifs are presented by both forward and reverse sequence scripts to simplify comparison and search.

**Fig. 1.**
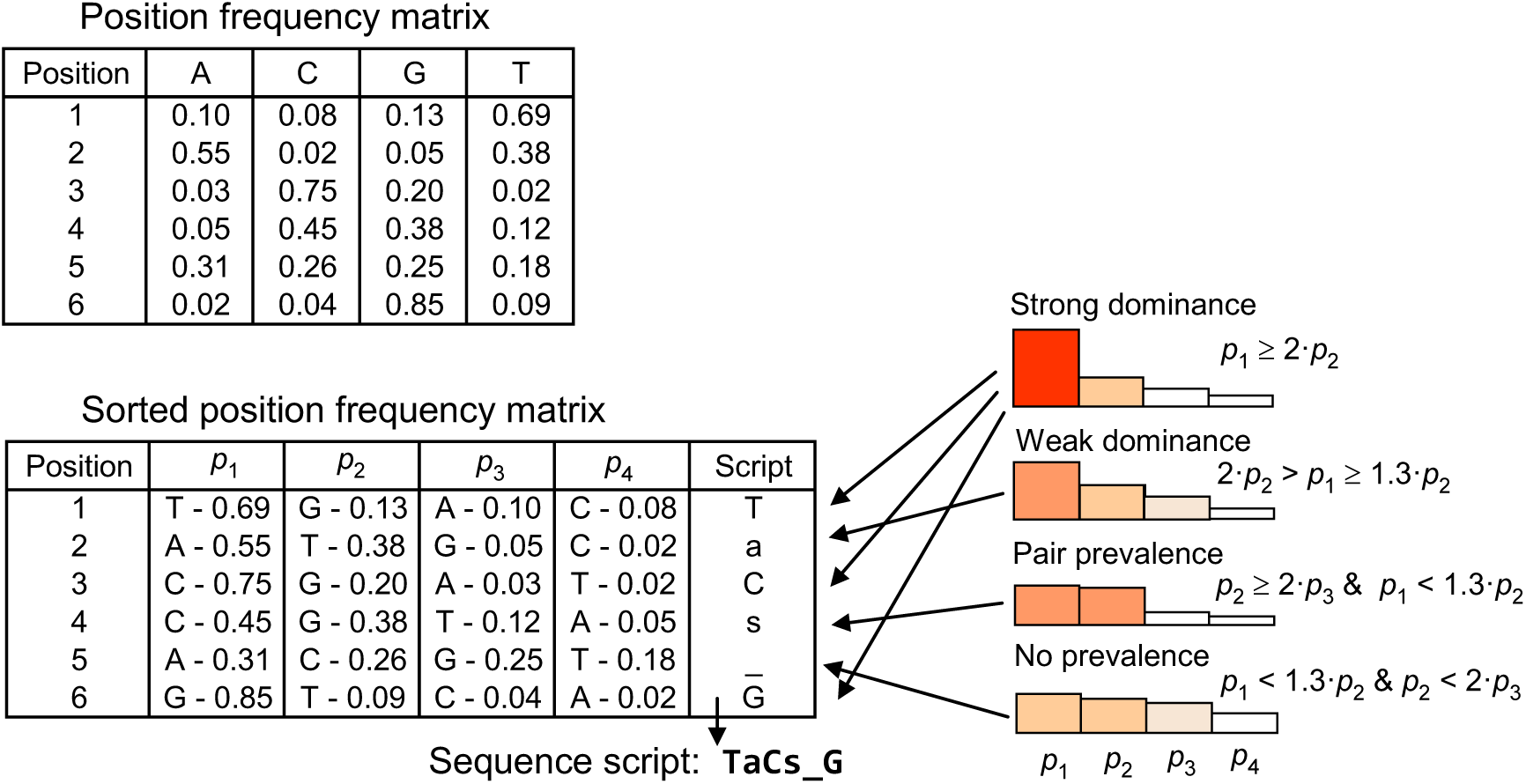
Making sequence script from a position weight matrix. In each position, nucleotides are sorted in the order of decreasing frequency, where *p_i_* is the frequency of a nucleotiode with rank *i*. The nucleotide symbol is entered as upper case if there is a strong dominance or lower case if there is weak dominance. Pair prevalence is marked by lower-case symbol (e.g., “s” stands for C/G), and no prevalence is marked by underscore.

## 3. Results

### 3.1. Generating a Compendium of DNA-Binding Motifs from ChIP-seq Data

The goal of this project is to generate reproducible DNA-binding motifs for most ChIP-seq data in the GEO database and establish all-against-all relations between TFs and DNA binding motifs as proposed by Ambrosini et al. (2020). Human and mouse ChIP-seq experiments were selected following information from the ENCODE project table (https://genome.ucsc.edu/encode/) and by additional search for individual TFs in GEO. Binding motifs of TFs were identified by reanalysis of 10,935 human and mouse ChIP-seq experiments (Supplementary file S1). DNA motifs enriched in ChIP-seq peaks were identified using CisFinder (Sharov and Ko 2009), and best 5 motifs from each experiment (if available) were selected for clustering. In addition, I used modified methods of motif search, where ChIP-seq peaks were filtered based on localization in promoters, or enhancers, or repeat-depleted regions (see Methods).

To cluster motifs, I developed a new k-mean clustering algorithm tailored for this task (see Methods). In comparison to hierarchical clustering, this approach is less computationally intensive and handles large data sets (e.g., 30,000 motifs and 2,000 clusters). After clustering, this software generates sequence strings (see Methods) which are aligned to each other within each cluster. The results are then loaded into Excel for visual inspection and manual post-processing, which includes merging similar clusters, splitting slightly heterogeneous big clusters, and removing outlier and redundant motifs.

Post-clustering QC was necessary to identify and remove low-quality ChIP-seq samples. A consistency score of a sample was estimated as the number of other samples targeting the same TF (or similar TFs^3^) that have matching motifs in the same cluster, summed up over all clusters where motifs of the tested sample are found. If consistency score was less than 1/10 of the maximum consistency score among all samples targeting the same TF, then the sample was considered low quality and is not reported. Low quality samples were more abundant in some individual GEO data sets, such as GSE151287 (Lai et al. 2021), where a new PRCP method for generating antibodies was tested. Thus, unknown motif clusters originating exclusively from such data sets were considered suspicious and were removed. In addition, some TFs have suspicious alternative binding motifs that show no resemblance to the main binding motif reported in the literature and databases. In such cases, I checked if the same samples yielded the main and alternative binding motifs. If samples in the alternative motif cluster failed to generate the main binding motif of the target TF, then the cluster was discarded as an artifact.

This compendium of DNA-binding motifs, which is called X-TFBS (“X” stands for all-versus-all cross comparison), includes 25,461 human motifs and 10,398 mouse motifs after filtering (Supplementary files S2 and S3). These motifs are derived from ChIP-seq experiments with 1497 human TFs and 624 mouse TFs^4^. These TFs included 459 human-mouse orthologous pairs (Fig. 2). The combined number of explored human and mouse TFs excluding orthologous duplicates is N = 1662, which is greater than the number of TFs in the HOCOMOCO database, N = 1577 (Vorontsov et al. 2024).

**Fig. 2.**
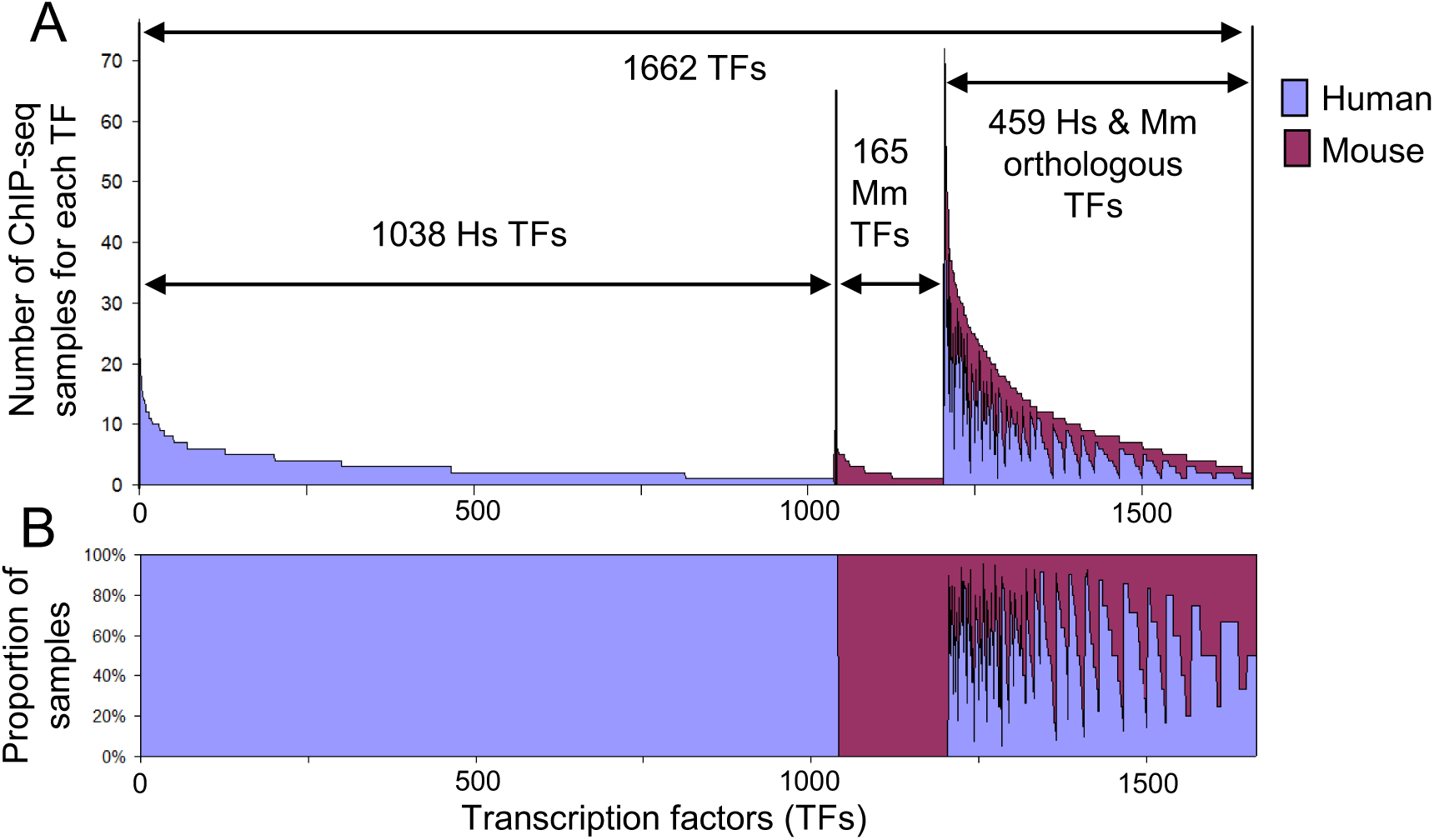
Composition of TFs and ChIP-seq samples, from which motifs in the X-TFBS compendium were derived. (A) Number of ChIP-seq samples; (B) proportion of human and mouse samples.

DNA-binding motifs in X-TFBS are grouped into 1184 clusters for human TFs and 563 clusters for mouse TFs (Supplementary files S4 and S5). These include 27 human and 26 mouse nonspecific clusters, which are not clearly associated with a particular TF. Cluster motifs that represent a consensus of three top regular motifs in each cluster are available in Supplementary files S6 and S7. Comparison of human and mouse clusters of motifs, excluding nonspecific ones, shows that 447 human motif clusters match fully or partially to mouse motifs, and 372 mouse motif clusters matched fully or partially to human motifs (Fig. 3A). A large portion of human-only and mouse-only motif clusters were identified from human-only and mouse-only TFs, respectively: 570 motif clusters (80.3% out of 710 human-only clusters) were derived from human-only TF’s ChIP-seq data, and 72 motif clusters (43.6% out of 165 mouse-only clusters) were derived from mouse-only TF’s ChIP-seq data.

**Fig. 3.**
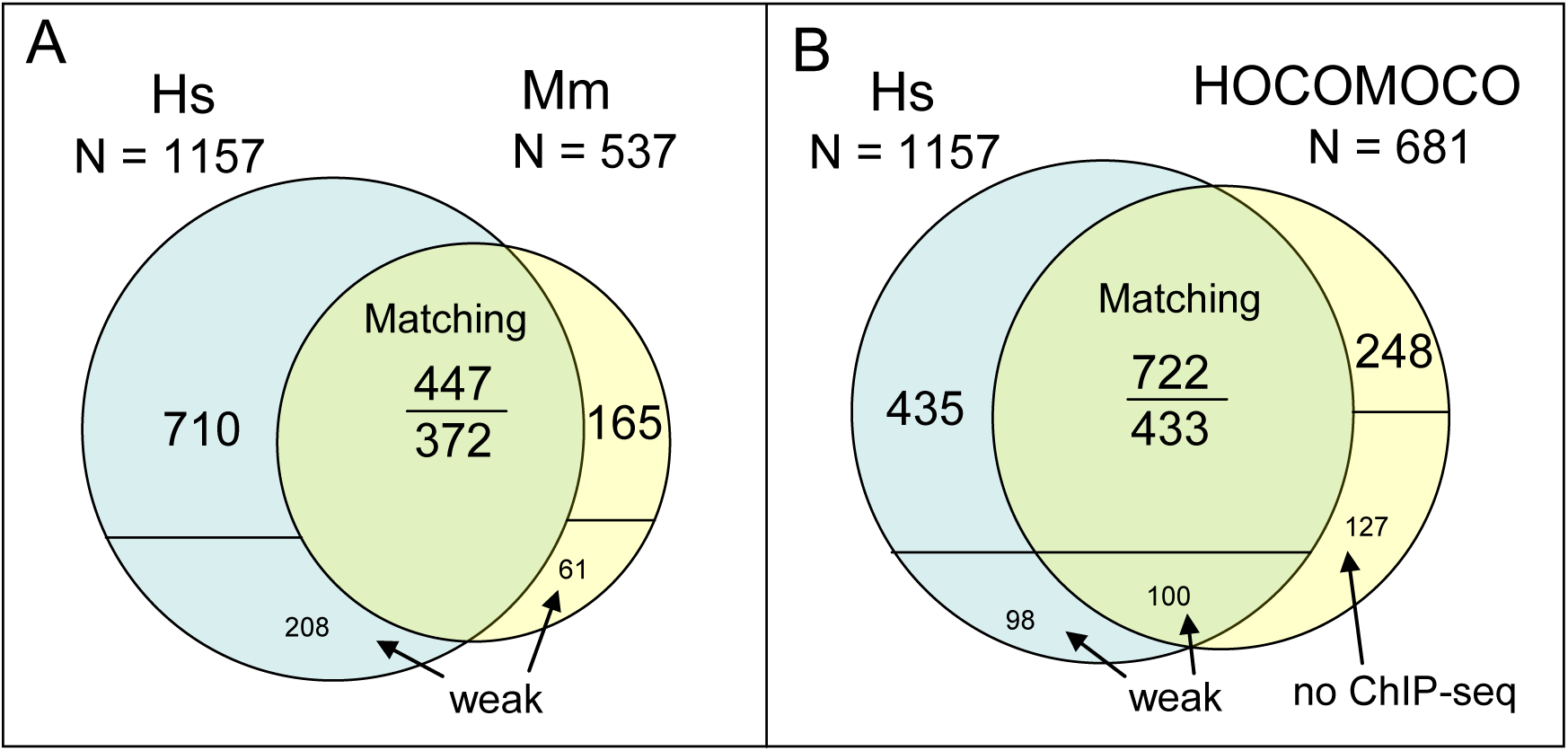
Relations between motif clusters: (A) in human (Hs) and mouse (Mm), and (B) in human and in HOCOMOCO database (Vorontsov et al. 2024). Overlap of motif clusters is based on similarity and quantified from the right set by a number above the line and from the left set by the number below the line. Weak clusters are those that are supported by a single RNA-seq sample.

Strongly similar DNA-binding motifs are found in 284 human-mouse pairs of clusters (Supplementary file S8), and most of them (N= 258) represent either orthologous pairs or TFs from the same gene family. Selected 114 human-mouse motif pairs together with motifs from published databases are shown in Table 1. In cases, where nonrelated human and mouse TFs bind to the same motif, one of the TFs is likely to bind indirectly. For example, ChIP-seq data for ZBTB33 is available for human but not for mouse, and its binding motif matches to the motif of its cofactor BRCA1 in both, human and mouse. Interaction of these two TFs bound to DNA is well characterized (Luo et al. 2025). Similarly, NANOG-SMAD1 cluster motif in human matches to the SOX2-repeat in mouse and SOX2 is a known cofactor of NANOG (Boyer et al. 2005).

**Table 1.**
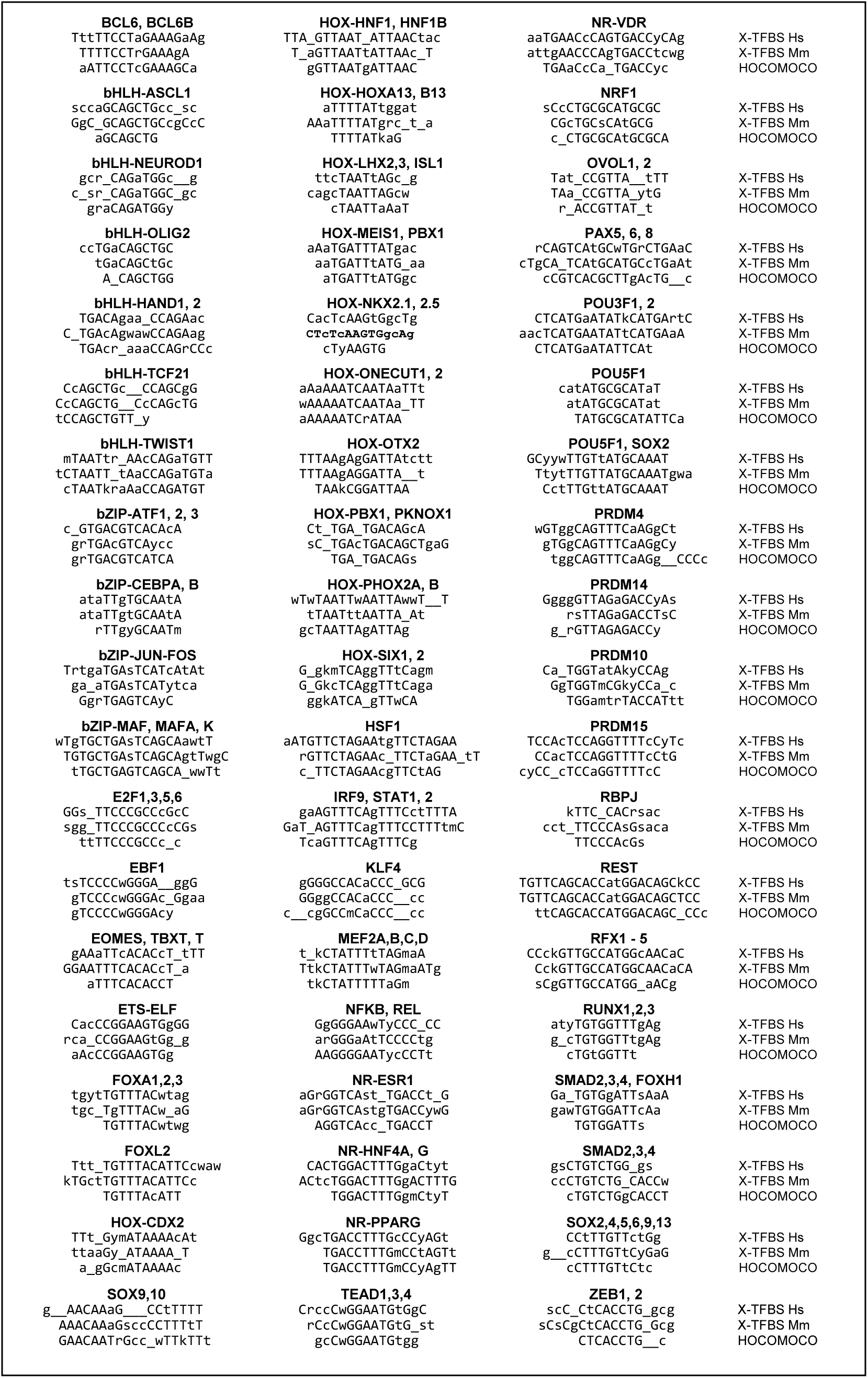
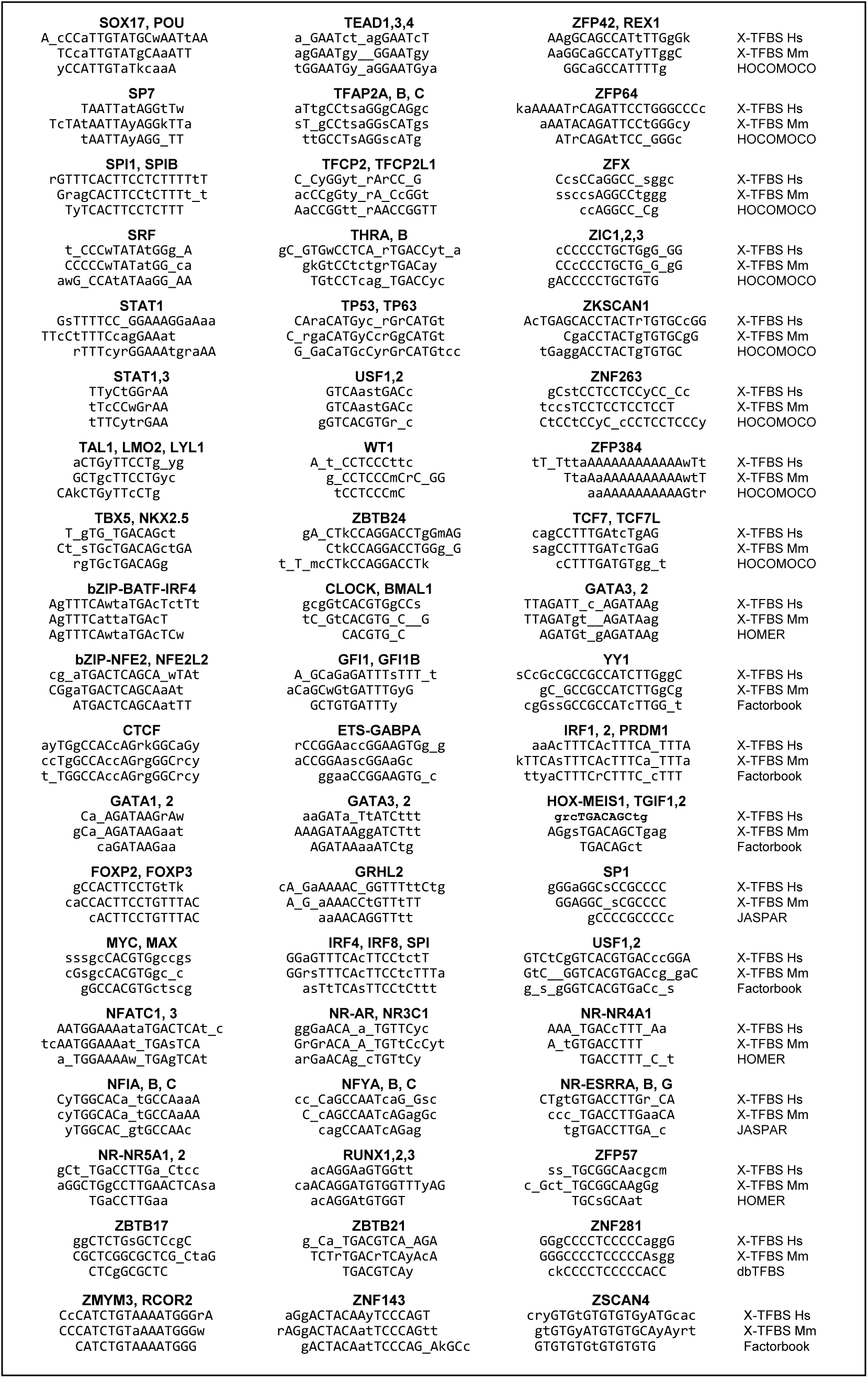
Selected 114 similar human (Hs) - mouse (Mm) motif pairs in X-TFBS.

Correspondence between TFs and motif clusters to which they bind is presented by pie charts (Fig. 4) and Supplementary file S9. TF-motif links are counted here if they are supported by not less than 25% ChIP-seq samples from the number of samples of the most strongly associated motif cluster, and by not less than 2 samples. In addition, TFs, whose names are included in the name of the motif cluster, are counted even if they do not match these conditions. These pie charts clearly show that the relation of TFs to binding motifs belongs to the “many-to-many” kind.

**Fig. 4.**
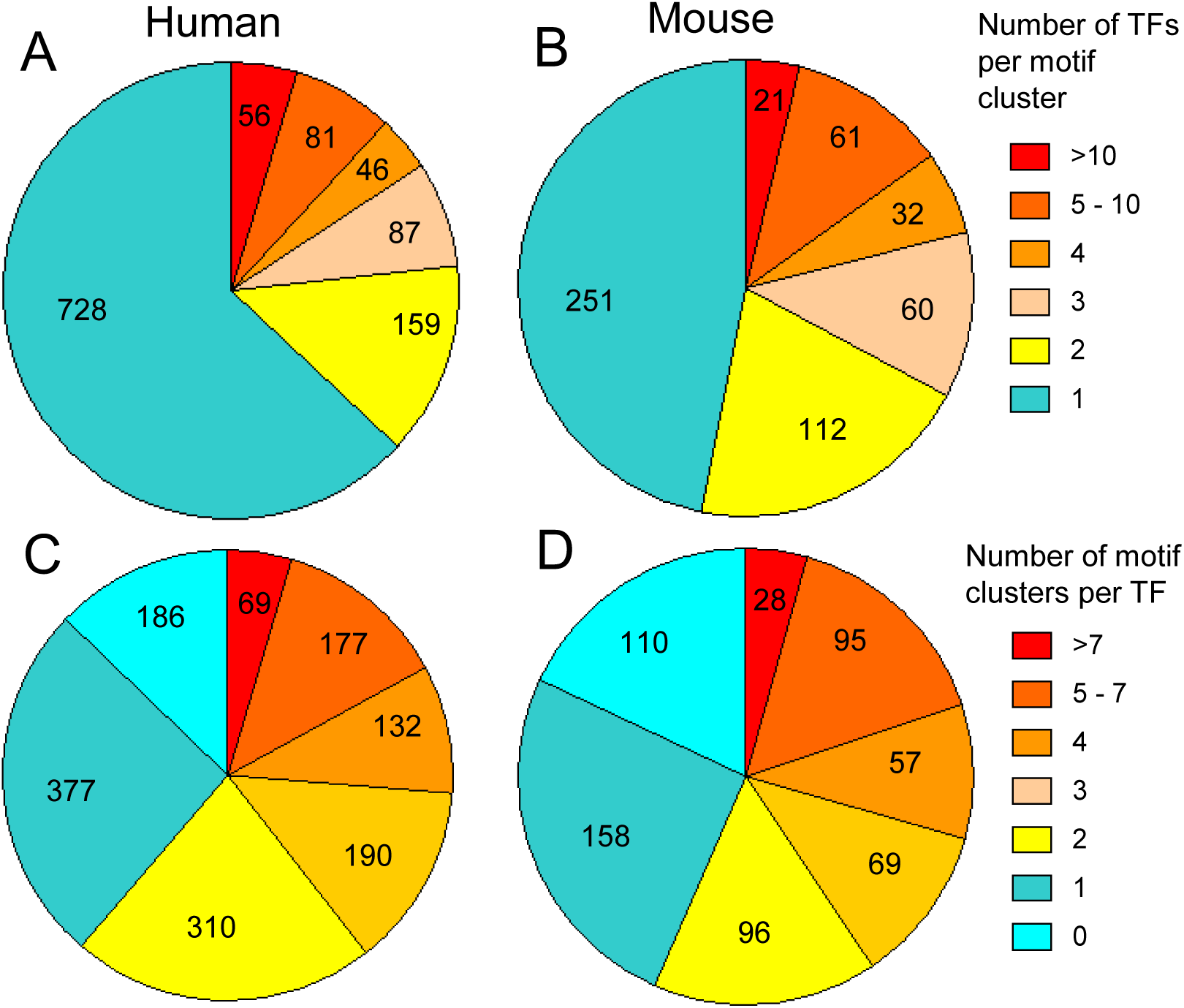
Relationship between TFs and their binding motifs in human (A,C) and mouse (B,D). A, B pie charts show the number of TFs that bind a motif cluster, and C, D pie charts show the number of distinct motifs bound by each TF.

About one third of human motifs and more than half of mouse motifs are bound by multiple TFs, and the proportion of TFs with a single known binding motif is just slightly above one quarter. About one fifth of TFs in human and mouse have no link to any motifs due to various factors. Some of them are not real TFs but chromatin modifiers or cofactors. The relation between TFs and motifs is not symmetric: more than a half of motifs in human and ca. half of motifs in mouse correspond to a single TF (Fig. 4A, B), whereas the proportion of motifs bound by a single TF is only ca. one quarter. This indicates that many TFs have alternative binding motifs, and many of these motifs are TF-specific (Fig. 4C, D).

### 3.2. Genome location of different motif clusters

To check if filtering of binding sites based on their location in promoters, enhancers, and repeat-depleted regions yields additional DNA-binding motifs of TFs, I compared motifs identified from all ChIP-seq peaks without filtering with clusters of X-TFBS compendium. It appeared that these motifs matched 852 clusters of X-TFBS for human and 421 clusters of X-TFBS for mouse. These cluster counts are by 26.4% and 21.5% smaller than the total counts of clusters for human and mouse, respectively, which proves that filtering of binding sites based on their location in promoters, enhancers, and repeat-depleted regions yields additional DNA-binding motifs of TFs.

Motifs clusters with individual motifs identified from ChIP-seq peaks located exclusively in promoters or enhancers were marked as “promoter” or “enhancer”, respectively. Other motif clusters were marked as “repeat” if they were detected from non-filtered ChIP-seq peaks, or from ChIP-seq peaks filtered as located in repeat-depleted regions (<70% repeats) without excluding motifs in repeats. There are relatively few promoter and enhancer motif clusters were infrequent: 1.1-1.7% and 4.1-4.5%, respectively, but repeat motif clusters are numerous: 38.6% in human and 23.7% in mouse (Fig. 5, upper charts).

**Fig. 5.**
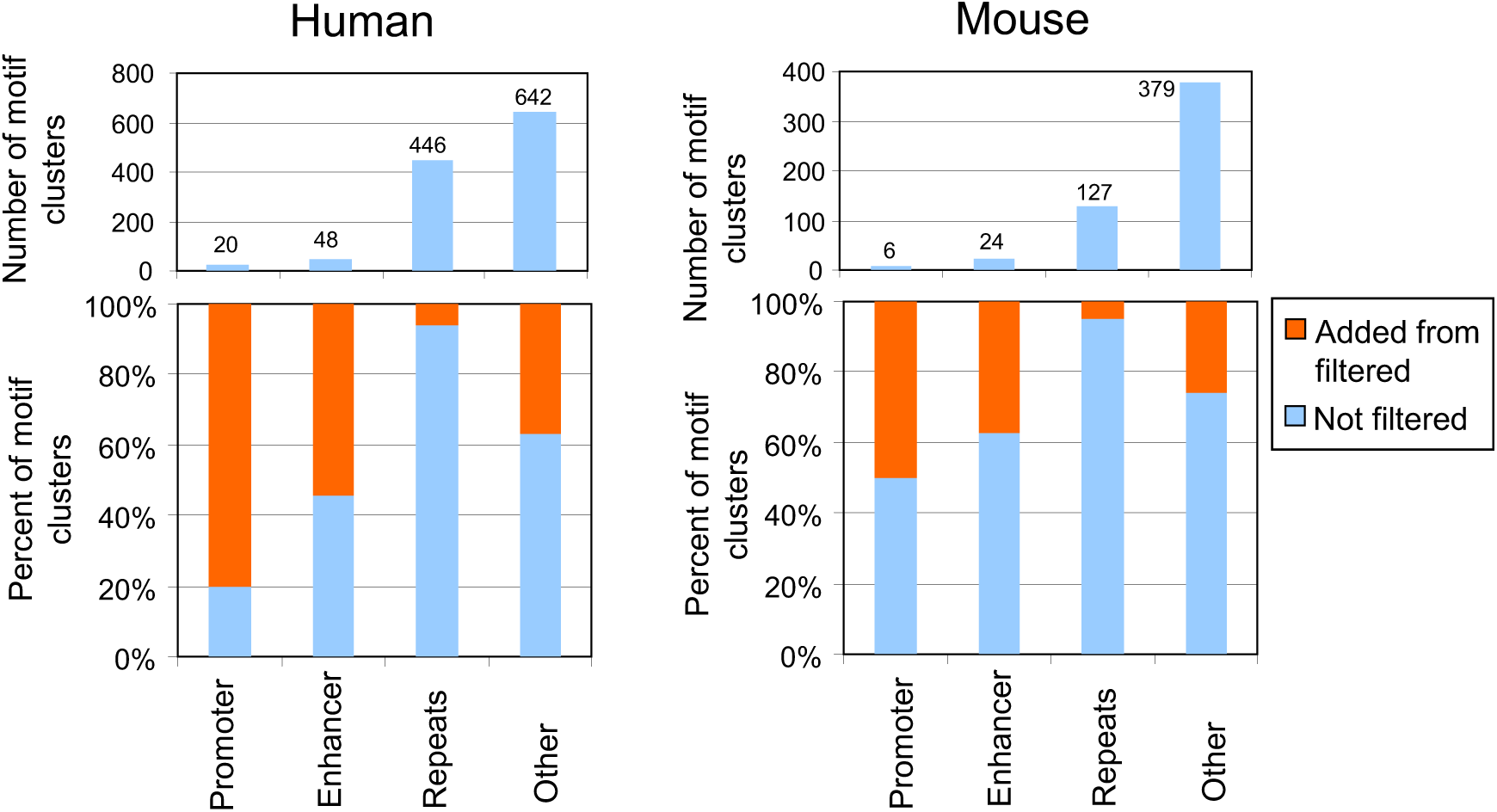
Relative abundance of motif clusters classified as “promoter”, “enhancer”, and “repeat” (upper charts) and the proportion of these motif clusters detected from non-filtered ChIP-seq peaks and additional motif clusters identified with location-specific filtering (lower charts) in human and mouse.

The majority of repeat motif clusters were detected from non-filtered ChIP-seq data, except a few motif clusters that were identified from repeat-depleted peaks (Fig. 5, lower charts). The largest number of additional motif clusters identified using filtered ChIP-seq peaks was in the category of “other” (i.e, neither promoter, nor enhancer, nor repeat), and these motifs were identified in ChIP-seq peaks with depleted repeats.

Genome coordinates of top three motifs from each cluster in X-TFBS were identified by searching corresponding ChIP-seq peaks for match to the PFMs “Search motifs” command in CisFinder. The numbers of found motif instances are shown in Supplementary files S4 and S5. The percent of motif instances in gene promoters (within 300 bp on both sides from TSS) was estimated for each motif cluster based on three top motifs (Supplementary file 10). The percent of motif matches in gene repeats was estimated as one minus ratio of motif matches in repeat-free genome regions within ChIP-seq peaks to motif matches in full length (200 bp) peaks for top three motifs. A scatterplot of percent of motifs in repeats versus their percent in promoters shows 4 subsets of motif clusters according to their localization in the genome (Fig. 6).

**Fig. 6.**
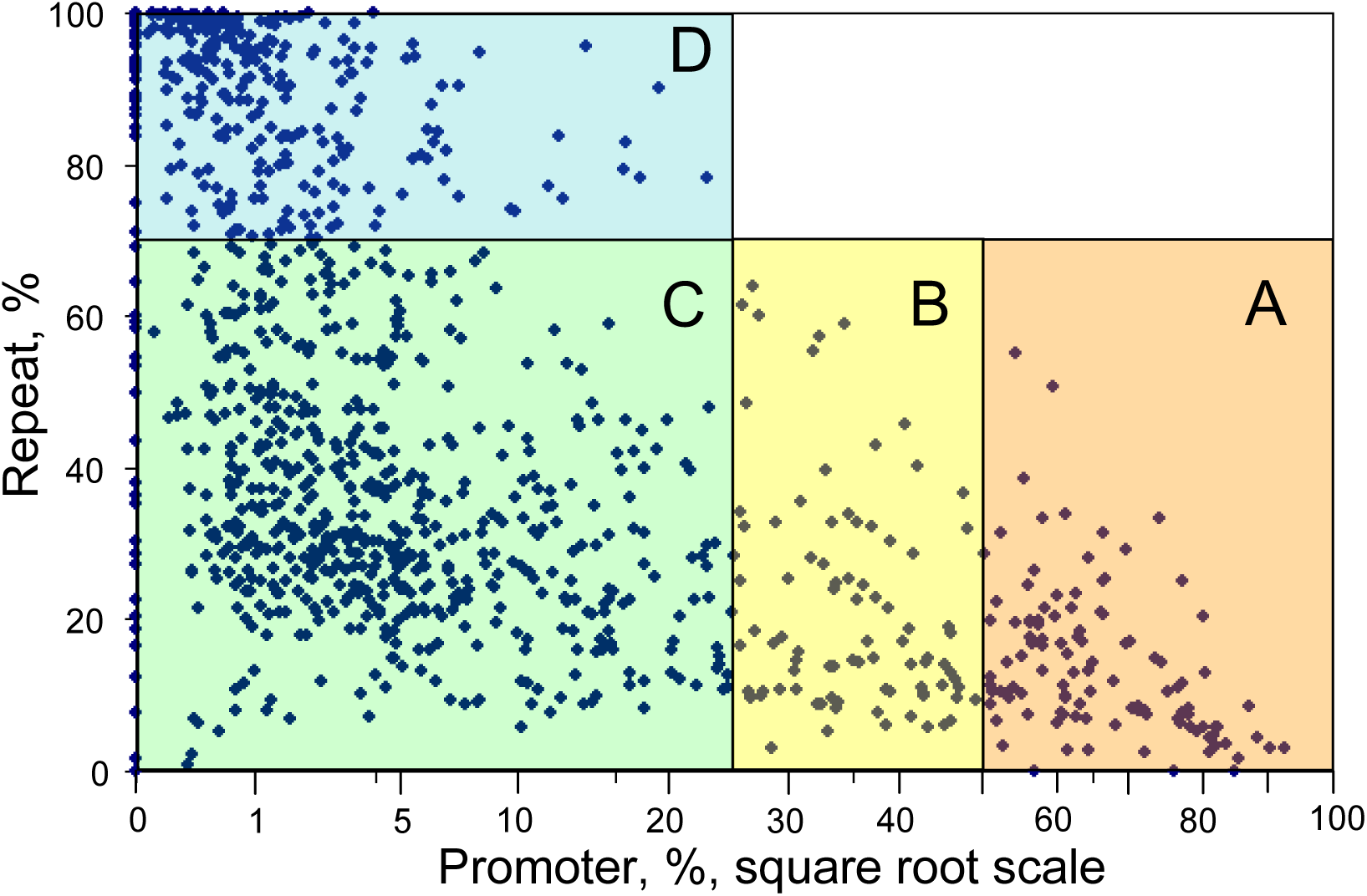
Four groups of human motif clusters that differ in their genome localization. Group A motifs are found mostly in promoters and in non-repeat DNA; group B motifs are present both in promoters and enhancers; groups C, and D of motifs are located in distal enhancers and have either low-to-moderate (C), or high-(D) percent located in repeats.

Each group of motif clusters is characterized by specific TFs that bind to these motifs, whereas only a few TFs have motifs that belong to multiple groups (Fig. 7). Details on TF specificity to groups of motif clusters is presented in Supplementary file 10. TFs specific to group A include many well-know promoter-bound factors that regulate recruitment of POL2 (TBP), mRNA elongation (NRLFE), cell cycle (E2F, MAX, MYCN, DMTF1), circadian rhythm (CLOCK, BMAL1, CRY, NPAS2, BHLHE40), gene repression (ETS-family TFs, YY1), activation of metabolic genes (NRF1, CREB1, CREM, USF1), hypoxia response (HIF1A), and embryo development (HES, HEY, KLF, OLIG2, USF2). TFs specific to group B have a strong association with immune response (NFKB, REL, MYB, STAT6, RFX, SOX9) and embryo development (GLIS1, POU3F1, POU3F2, TFAP2, WT1, PKNOX1). Group C of motif clusters is associated with the largest number of TFs involved in various functions, such as embryo development (ASCL1, TCF12, ATOH1, MYOD1, NEUROG2, CDX2, EOMES, TBXT, TBX3, HAND2, SMAD1, SMAD3, POU5F1, SOX2, NANOG, SOX17), immune response (BCL6, BCL6B, IKZF1, IRF1, IRF4, IRF8, SPI1, SPIB, MAF, MAFK, RUNX1, STAT3, TP53), hormone metabolism (AR, ESRR, NR3C1, PGR, HNF4, PPARG, RXR, RAR), transcription repression (REST, CTCF). Zinc finger C2H2 proteins dominate among TFs specific to the group D of motif clusters, which are located mostly in genomic repeats. It is assumed that their main function is repression of transposons (Imbeault et al. 2017).

**Fig. 7.**
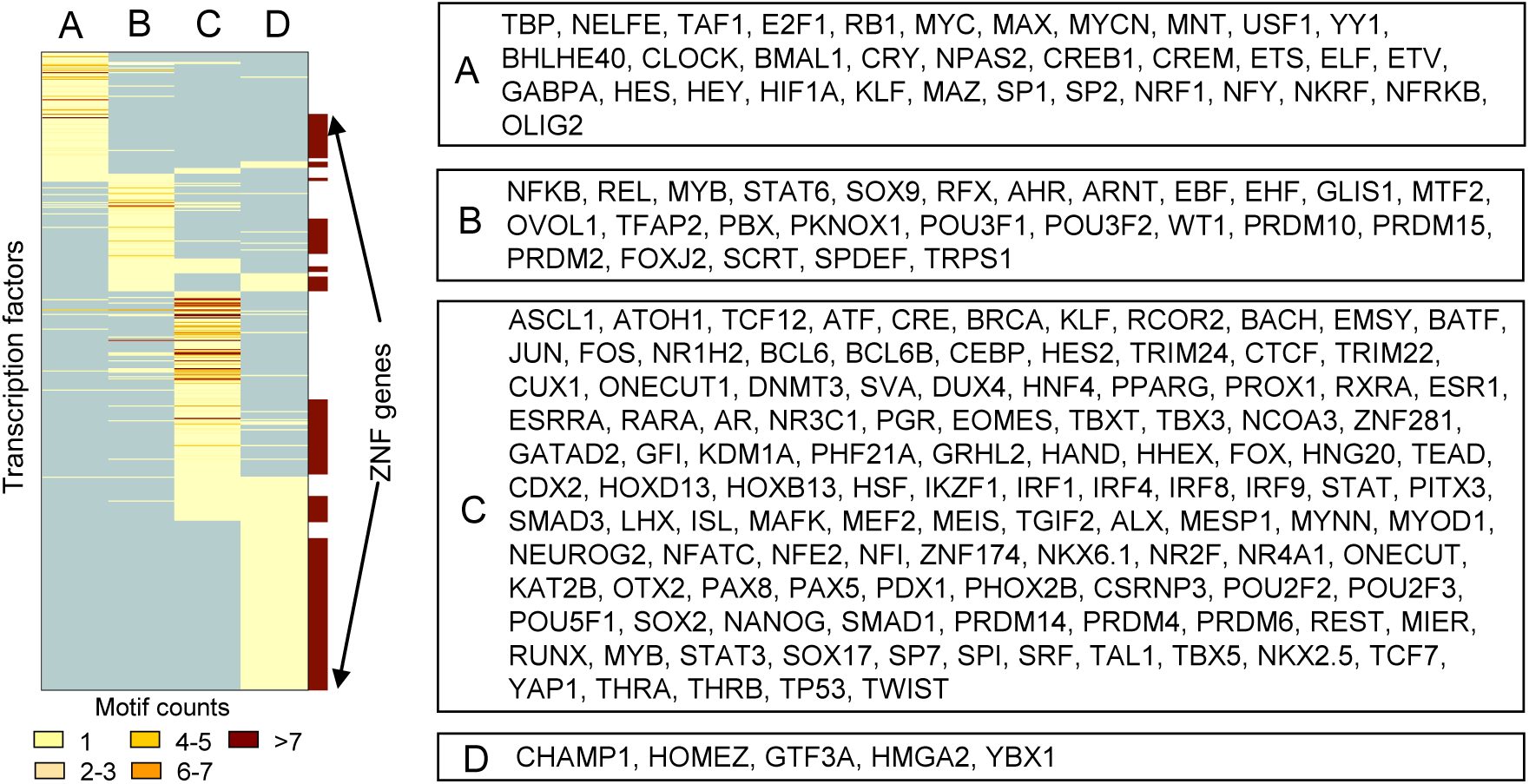
TFs that bind motif clusters in groups A, B, C, and D (see Fig. 6). Zinc finger ZNF genes are marked at the right side of the heatmap (symbols are not shown).

### 3.3. Comparison of X-TFBS DNA-binding motifs with other databases

To evaluate the relative scope of the X-TFBS compendium, I applied the k-mean clustering method to motifs in existing databases of motifs. The number of motif clusters was 682 in HOCOMOCO (Vorontsov et al. 2024), 294 – in Factorbook (Pratt et al. 2022), 267 – in the ChIP-seq portion of University of Toronto database (Lambert et al. 2018), 193 – in HOMER (Heinz et al. 2010), and 336 – in the combined data from dbTFBS (Yu et al. 2021), JASPAR (Rauluseviciute et al. 2024), and SwissRegulon (Pachkov et al. 2013). The combined number of human motif clusters and mouse-only motif clusters in the X-TFBS compendium is 1322, which makes it the largest among databases of DNA-binding motifs.

The majority of DNA-binding motifs in the X-TFBS compendium match to already known motifs. Out of 1157 human motifs, 722 (62.4%) match fully or partially to 433 HOCOMOCO motifs (Fig. 3B). However, some motifs reported in the public databases are missing in the X-TFBS compendium. For example, HOCOMOCO has 248 (36.4%) motif clusters that do not match to human motifs in X-TFBS (Fig. 3B). Out of these, a large subset of motifs (N = 127) comes from in-vitro SELEX and PBM-array experiments, which may generate erroneous motifs because they do not replicate in vivo conditions. Examples of erroneous DNA-binding motifs in HOCOMOCO that are supported by multiple data sets are shown in Supplementary file S11. Other 121 human motif clusters from HOCOMOCO that are not present in X-TFBS include the following: (i) 24 motifs of TFs that are not processed in this study (e.g., ATMIN, CPXCR1, DNTTIP1, GTF2IRD2, MYRFL, NPAS4, SP100, and TET3); (b) 15 motifs that are modifications of motifs present in X-TFBS (e.g., motif PTF1A in HOCOMOCO is a combination of the core PTF1A and RBPJ^5^ motifs, which are both present in X-TFBS); and (c) 3 of them match to mouse motifs in X-TFBS. Factorbook and University of Toronto databases have 20 and 29 motif clusters, respectively, that do not match to X-TFBS.

The X-TFBS compendium includes 300 and 145 new motif clusters^6^ for human and mouse, respectively (Table 2). These new motifs comprise ca. 26% of all human and mouse motif clusters. More than a third of all new motifs are weak (i.e., supported by a single ChIP-seq sample) and may include more artifacts than other motifs supported by multiple experiments. A large portion of new motif clusters belong to C2H2 zinc finger TFs: 215 in human clusters (71.43%) and 49 in mouse (36.96%). This group of fast-evolving TFs requires more extensive ChIP-seq studies because half of new C2H2 zinc finger motifs are weak (N = 100 for human and N = 31 for mouse). Another large group of new DNA-binding motifs belong to secondary motifs: 130 human motifs (43.19%) and 64 mouse motifs (46.38%). Most of these motifs are modifications of the main motif pattern via alteration of flanking regions or rearrangement of binding components (e.g., homodimers, heterodimers, or palindromes). Summarizing these data, most new motifs are for C2H2 zinc finger TFs, and almost half of new motifs are secondary ones.

**Table 2.** Numbers of clusters of TF DNA-binding motifs.

| Clusters of DNA-binding motifs | Human |  | Mouse |  |
| --- | --- | --- | --- | --- |
|  | N | % vs Total | N | % vs Total |
| All motif clusters | 1157 | 100.00% | 537 | 100.00% |
| Match published motifs | 858 | 74.16% | 391 | 72.81% |
| New motifs | 299 | 25.84% | 146 | 27.19% |
|  | N | % vs New | N | % vs New |
| New motifs weak | 105 | 35.12% | 51 | 34.93% |
| New motifs zinc fingers | 215 | 71.91% | 49 | 33.56% |
| New secondary motifs | 129 | 43.14% | 64 | 43.84% |

### 3.4. Shared binding motifs indicate potential interaction of TFs

Sharing motifs between different TFs is a wide-spread phenomenon (Fig 4A,B), which points to potential functional interactions between these TFs. Here I present examples of such interactions supported by published research.

1. In human, TBX5 and NKX2-5, which are key factors regulating heart morphogenesis, both bind to motifs T_gTG_TGACAGct or GgTGTTGACAGc^7^, which are TBX-specific. Where NKX2-5 binds alone, its motif is NKX2-specific: t_ttgAGTG_ctG_a or cAgcCACTTgAgA. In mouse, Nkx2-5 binds to the NKX2-specific motifs: cTtGAGTGgc or cTgcCACTTGAg_g, but it does not bind to the Tbx5-Tbx20 motif CT_sTGcTGACAGcwGA. This is an example, where sharing a binding motif between TFs exists in human but not in mouse. Direct molecular interaction of human TBX5 and NKX2-5 has been confirmed by X-ray crystallography (Pradhan et al. 2016).
2. Three TFs TAL1, LMO2, and LYL1 that are known to support branching angiogenesis (Yamada et al. 2022) and hematopoiesis (Hoang et al. 2016) bind to the same motif aCTGyTTCCTg_yg, which is a shorter version of a TAL1-specific motif cwsCAGCTGyTTCCTG_CG. These proteins bind together and collectively activate hematopoietic genes (Hoang, Lambert and Martin 2016). TAL1 also binds to GATA motifs (Ca_AGATAAGrAw and TC AGATAAGA_aa) and it is known to interact with GATA2 to initiate vessel spouting (Yamada et al. 2022), and with GATA3 (Sanda et al. 2012).
3. ARID3A is known to repress binding and activity of TFs HNF4A and HNF1A in regulating intestinal epithelial proliferation and differentiation (Angelis et al. 2024). It appears that ARID3A and HNF4 share nearly the same motif: tTGrACTTTGaac and CGCTGgACTTTGg_Cyyc, respectively, which indicates potential direct interactions between these TFs. Another motif of ARID3A is AAtyATTAACwA matches the motif TTA_GTTAAT_ATTAACtac of HNF1A and HNF1B, which is also a sign of potential interaction.
4. SMAD1 and SMAD4 bind to a CG-rich motif CssCTGGCGCC_gcgg, which is similar to motif CTGGaGCC reported in HUVEC cells stimulated by BMP9 (Morikawa et al. 2011). Accorting to (Massague et al. 2005), SMAD1 and SMAD4 bind to DNA as a dimmer. However, SMAD1 has several other binding partners. In embryonic stem cells it binds together with POU5F1, SOX2, and NANOG to motif GCyywTTGTtATGCAAAT (X-TFBS), which is consistent with earlier findings (Chen et al. 2008). According to X-TFBS, SMAD1 also binds to motif CaccCtGGAATGtGgc (cluster TEAD-SMAD1-GATAD2-YAP1). According to (Yao et al. 2014), phosphorylated SMAD1-SMAD4 complex binds to YAP1 and disrupts TEAD1-YAP1 interaction, which results in the repression of proliferation of mouse neural stem cells.
5. SMAD2 and SMAD3 have entirely different DNA-binding motifs than SMAD1 thanks to their interaction with FOXH1, which binds to motif Ga_TGTGgATTsAaA and three alternative motifs gsCTGTCTGG_gs, g_AGTCTG_CACCTTy, and tGTCTG_CACCtc (X-TFBS). The ending part of the latter two motifs matches to the binding motif of EOMES, which is a known cofactor of FOXH1 (Slagle et al. 2011). SMAD2 and SMAD3 get phosphorylated in response to Nodal TGF-β signals (Aragon et al. 2019). After that, they form hetero-trimeric complexes with SMAD4 (Miyazono et al. 2018), enter the nucleus where they bind to FOXH1 by displacing Tle/Groucho cofactor (Charney et al. 2017).
6. TFs from the E-box motif cluster gcgGtCACGTGgCCs that includes CLOCK, BMAL1, CRY1, CRY2, NPAS2, MAX in human are all involved in the circadian clock function (Blazevits et al. 2020; Fahrenkrug et al. 2008; Landgraf et al. 2016; Tamayo et al. 2015) and interact at these binding sites. In particular, CLOCK forms heterodimers with BMAL1, which bind to E-box and activate the expression of CRY1, CRY2, PER1, and PER2 genes. As CRY and PER proteins accumulate in the cell, they form oligomers and enter the nucleus of cells, where they repress activating functions of CLOCK and BMAL1 (Albrecht 2012; Hirota and Fukada 2004; Mohawk et al. 2012). ChIP-seq data on PER1 and PER2 in human was not found, but *Per1* and *Per2* in mouse appear to bind a similar circadian E-box motif tC_GtCACGTG_C G (X-TFBS). Other TFs in human and mouse circadian motif clusters include ESR1, APC, CSRNP1, EEA1, NR2E3. ESR1 is a circadian regulator (Royston et al. 2014); APC, CSRNP1, and EEA1 are circadian-regulated TFs (Carvalho Cabral et al. 2024; Hattar et al. 2002; Maissan and Carlberg 2025), and NR2E3 is a cofactor of a circadian clock TF NR1D1 (Cruz et al. 2014). BHLHE40 is another circadian gene (Kato et al. 2014), which has a similar E-box motif sCGGkCACGTGmcCs in human and Tc_GTCACGTGacca in mouse.
7. Based on X-TFBS, insulator and chromatin loop organizer CTCF has multiple motif clusters, and some of them are shared with cohesins RAD21, SMC3, and STAG1, and other TFs, such as ZNF384 and TRIM22. Interaction of CTCF with these proteins has been also identified from experiments (Choi and Kim 2025; Ibn-Salem and Andrade-Navarro 2019; Kuang and Wang 2020; Rubio et al. 2008). The main motif cluster with combined CTCF and RAD51 binding is sgCCACcAGrgGGCaGyg, which is slightly different from the canonical CTCF motif cTGgCCACCAGgGGGCGC (X-TFBS).
8. E2F1, E2F3, E2F4, E2F5, E2F6, RB1, TFDP1, TFDP2, and LIN9 bind to the same motif cluster GGs_TTCCCGCCcGcC. In addition, E2F7, E2F8, and RB1 bind to motif cluster sG_tTTCCCGCCAaAa. These proteins are members of the DREAM complex that regulates cell proliferation and apoptosis (Cao et al. 2010; Ding et al. 2018; Forristal et al. 2014; Tian et al. 2007).
9. GABPA and ZBTB11 bind to the same cluster motif gc_gCCGGAAGTGgcG. According to (Wilson et al. 2020), ZBTB11 interacts with GABPA to regulate mitochondrial functions.
10. The main binding motif of the homeodomain TF MEIS1 is aAaTGATTTATgac, and several other TFs bind to the same motif, such as PBX2, PKNOX1, HOXA9, and HOXA10. It appears that MEIS1, PBX2, and HOXA9 form triple complexes at such binding sites in myeloid leukemia cells (Shen et al. 1999). Interaction of PBX2 and HOX10 was observed in the endometrium, and expression MEIS1 and HOX10 was menstrual cycle specific (Sarno et al. 2005); authors hypothesize the existence of trimeric complexes MEIS1-PBX2-HOX10. Interaction of MEIS1 with PKNOX1 may require PBX protein (Knoepfler et al. 1997; Mian and Zeleznik-Le 2016). Another binding motif grCTGACAGCTg is shared by MEIS1 and TFIG2. Although interaction of MEIS1 with TGIF2 has not been observed, it was found that TGIF1 interferes with a MEIS1-dependent transcription regulation by associating with MEIS1-bound genome locations (Willer et al. 2014).
11. Immune response regulators IRF1 and IRF2 bind to the same DNA motif aaAcTTTCAcTTTCA_TTTA, which is also shared with a repressor PRDM1. It has been shown that PRDM1 binds to the same sites as IRF1 and interferes with IRF1-dependent transcription activation after interferon-gamma (IFN-γ) stimulation (Mould et al. 2015). IRF2 is considered as a competitor of ITF1 for binding to their common motif because IRF2 reduces the action of IRF1 (Hochhaus et al. 1997).
12. IRF4 and IRF8 bind to the same motif GGaGTTTCAcTTCCtctT which is shared with SPI1, ETV6, and IKZF1-IKZF2. IRF4, IRF8, and SPI1 are known to interact directly at their shared motif to regulate early development of B-cells and suppress leukemia (Pang et al. 2016). IRF8 was shown to recruit ETV6/TEL in an IFN-γ-dependent manner in macrophages and mediate repression of IFN-γ effect in a negative feedback way (Kuwata et al. 2002). Also, IRF4 and IRF8 are known to cooperate with IKZF1 (Collin and Bigley 2018).
13. IRF9 binds to motif a_aAGTTTCAgTTTCctTTTAc together with STAT1 and STAT2. These three TFs are known to bind together to form a ISGF3 complex responsible for processing of non-canonical IFN-γ signaling (Fink and Grandvaux 2014). In addition, IRF9 forms a complex with STAT2 to perform STAT1-independent signaling (Fink and Grandvaux 2014; Rengachari et al. 2018).
14. NFYA, NFYB, and NFYC bind to the same CCAAT motif cc_CaGCCAATcaG_Gsc, which is shared with many other TFs, including SP1 and SP2. Direct interaction of these TFs with SP1 and SP2 has been reported (Cui et al. 2023; Roder et al. 1999). In particular, activation of the E2F1 gene (and the cell cycle) is mediated by binding of a NFY-SP1 complex to the promoter of E2F1 after 17beta-Estradiol stimulation of MCF-7 cells (Wang et al. 1999). Because the CCAAT motif is strongly bound by E2F and TFDP1-TFDP2, I hypothesize that E2F-TFDP complex can repress NFY activity via protein-protein interaction in a negative feedback way.
15. Androgen receptor AR, glucocorticoid receptor NR3C1, and progesterone receptor PGR bind to the same motif GrGaACAga_TGTTCcc (+ two alternative similar motifs). It is not clear how these receptors have distinct transcriptional programs despite of the strong similarity of binding motifs (Kulik et al. 2021). Possibly, coactivators determine the interaction of proteins. For example, FOXA1 binding facilitates replacement of AR by NR3C1 (Helminen et al. 2023). Notably, FOXA1 binds this motif according to X-TFBS. Another potential coactivator of NR3C1 is AP-1 (JUN+FOS), which is shown to stimulate transcriptional activation of *Lhcgr* by NR3C1 (Zhang et al. 2022). In X-TFBS, weak AP-1 binding sites (motif cluster NONSPEC6 JUN-like) are bound by NR3C1 but not by AR or PGR.
16. Estrogen receptor ESR1 binds to motif aGrGGTCAst_TGACCt_G, which is also bound by FOXM1, RARA, RARG, and NCOA3. FOXM1 is a transcriptional target of ESR1 (Millour et al. 2010; Peng et al. 2025); thus, its co-binding to the activating TF site may provide a negative feedback effect. Retinoic acid receptors RARA and RARG tend to have cross-inhibiting effects with estrogen receptor (Jimenez-Dominguez et al. 2021). Thus, they most likely inhibit regulatory effects of ESR1. Finally, NCOA3 is a primary co-activator of ESR1 and binds it directly (Yi et al. 2017).
17. Transcriptional repressor REST binds to motif TGTTCAGCACCatGGACAGCkCC, and this motif is shared with repressor EHMT2, which is a H3K9 methyltransferase. EHMT2 interacts with REST via CDYL protein (Mulligan et al. 2008).
18. USF1, USF2, RAD51, and MITF have the same binding E-box motif GTCtCgGTCACGTGACccGGA. ChIP-seq peaks of USF1, USF2, MITF, and RAD51 largely overlap in several human cell lines, which points to potential functional interaction between them (Kang et al. 2021).
19. Transcriptional activator and repressor YY1 binds to motif sCcGcCGCCGCCATCTTGggC, and PHF8, CDK9, and SIN3A have the same binding motif based on ChIP-seq. These TFs interact as follows from the facts that PHF8 demethylates YY1 protein (Wu et al. 2024), CDK9 forms elongation complex with YY1 (Qiu et al. 2021), and SIN3A is a co-repressor of YY1 (Lu et al. 2011).
20. Palindromic motifs c_cTCTCGCGAGAg and CcAGCTCTCGCGAGa are enriched in ChIP-seq peaks of ZBTB33, BRCA1, BANP, and DYRK1A in human. In mouse, however, there is no ChIP-seq data on *Zbtb33* and *Brca1*, but *Banp* and *Dyrk1a* have a matching motif tCcArcTCTCGCGAGAgagG. It has been shown that BRCA1 is recruited to DNA-bound ZBTB33 (Luo, Tripathi, Liu, Chen, Cowan, Daniel, Yan, Richmond and Reynolds 2025).
21. ZNF143 binding motif is aGgACTACAAyTCCCAGT, and the same motif is enriched in ChIP-seq peaks of THAP11, HCFC1, RBPJ, and NOTCH1 in human, and motif rAGgACTACAatTCCCAGtt is enriched in ChIP-seq peaks of Znf143, Hcfc1, and Notch1 in mouse. According to (Vinckevicius et al. 2015), ZNF143 binds to the ACTACA submotif first and then directs the recruitment of THAP11 and HCFC1 to ZNF143-occupied loci. Another study indicates strong overlap of ZNF143 and NOTCH1 binding sites, and that RBPJ can replace ZNF143 in ZNF143-NOTCH1 complexes (Wang et al. 2011).
22. Bcl6 binding motif TTTCcAGGAAA (in mouse) is enriched in ChIP-seq peaks of Bcl6b, and Stat5a. Bcl6b is a known repressor of Bcl6; also Bcl6b and Bcl6 can bind to similar motifs of STAT TFs in a competitive way (Hartatik et al. 2001).

These examples indicated that all-against-all association of TFs and their DNA-binding motifs yields evidence of potential interaction between TFs located at their binding sites, and it may stimulate analysis of additional bio-molecular interactions in gene regulatory networks.

## Supporting information

List of ChIPseq samples

Human motifs (MEME)

Mouse motifs (MEME)

Clustered human motifs

Clustered mouse motifs

Human cluster motifs (MEME)

Mouse cluster motifs (MEME)

Human-mouse similar motif pairs

Relations of transcription factors to motif clusters

Frequency of human motifs in relation to promoters and repeats

Erroneous motifs derived from non-ChIPseq data

## Declarations

I declare no conflicting financial and non-financial interests in publishing this research. This project was not funded and not supported in any other way by any person or organization, including my employer Ricoh Biosciences, and I have completed it in my free time. No confidential materials are used in this project.

## Author approval

the author has seen and approved the manuscript, and it hasn’t been accepted or published elsewhere.

## List of supplementary files

Supplementary File S1. List of ChIP-seq samples: human and mouse

Supplementary File S2. Human ChIP-seq motifs, meme format

Supplementary File S3. Mouse ChIP-seq motifs, meme format

Supplementary File S4. Clustered human ChIP-seq motifs

Supplementary File S5. Clustered mouse ChIP-seq motifs

Supplementary File S6. Human cluster motifs, meme format

Supplementary File S7. Mouse cluster motifs, meme format

Supplementary File S8. Human and mouse matching motifs

Supplementary File S9. Correspondence between TFs and motif clusters

Supplementary File S10. Frequency of 3 top motifs in human motif clusters in relation to promoters and repeats

Supplementary File S11. Erroneous motifs in HOCOMOCO derived from non-ChIPseq data

## Footnotes

1 Code is available at https://kolab.elixirgensci.com/cisfinder/download.html and https://github.com/AlexeiSharovBaltimore/CisFinder

2 TSS – transcription start site.

3 Similar TFs are those that belong to the same protein family (e.g., ATF, KLF, SOX, FOX, NFI), easily identifiable by the first three characters of a gene symbol. Exceptions are zinc-finger TFs: ZNF, ZFP, PRDM, ZBTB, ZSCAN, ZBED, SP, where similar gene symbols do not imply similarity of binding motifs.

4 Besides TFs that directly bind DNA motifs, these sets of ChIP-seq targets include chromatin modifiers and other co-factors interacting with TFs.

5 RBPJ motif is supported by (Jin and Xiang 2019).

6 Here ‘new motifs’ means not found in public databases HOCOMOCO, Factorbook, HOMER, JASPAR, Toronto database, SwissRegulon, and dbTFBS. But some of these motifs may be present in published articles.

7 In this section I use Consolas font for DNA-binding motifs of TFs, and use bold face for the core portion of motifs.

